# Preterm birth alters the development of cortical microstructure and morphology at term-equivalent age

**DOI:** 10.1101/2021.06.03.446550

**Authors:** Ralica Dimitrova, Maximilian Pietsch, Judit Ciarrusta, Sean P. Fitzgibbon, Logan Z. J. Williams, Daan Christiaens, Lucilio Cordero-Grande, Dafnis Batalle, Antonios Makropoulos, Andreas Schuh, Anthony N. Price, Jana Hutter, Rui PAG Teixeira, Emer Hughes, Andrew Chew, Shona Falconer, Olivia Carney, Alexia Egloff, J-Donald Tournier, Grainne McAlonan, Mary A. Rutherford, Serena J. Counsell, Emma C. Robinson, Joseph V. Hajnal, Daniel Rueckert, A. David Edwards, Jonathan O’Muircheartaigh

## Abstract

**Introduction:** The dynamic nature and complexity of the cellular events that take place during the last trimester of pregnancy make the developing cortex particularly vulnerable to perturbations. Abrupt interruption to normal gestation can lead to significant deviations to many of these processes, resulting in atypical trajectory of cortical maturation in preterm birth survivors.

**Methods:** We sought to first map typical cortical micro and macrostructure development using *invivo* MRI in a large sample of healthy term-born infants scanned after birth (n=270). Then we offer a comprehensive characterisation of the cortical consequences of preterm birth in 78 preterm infants scanned at term-equivalent age (37-44 weeks postmenstrual age). We describe the group-average atypicality, the heterogeneity across individual preterm infants, and relate individual deviations from normative development to age at birth and neurodevelopment at 18 months.

**Results:** In the term-born neonatal brain, we observed regionally specific age-associated changes in cortical morphology and microstructure, including rapid surface expansion, cortical thickness increase, reduction in cortical anisotropy and increase in neurite orientation dispersion. By term-equivalent age, preterm infants had on average increased cortical tissue water content and reduced neurite density index in the posterior parts of the cortex, and greater cortical thickness anteriorly compared to term-born infants. While individual preterm infants were more likely to show extreme deviations (over 3.1 standard deviations) from normative cortical maturation compared to term-born infants, these extreme deviations were highly variable and showed very little spatial overlap between individuals. Measures of regional cortical development were associated with age at birth, but not with neurodevelopment at 18 months.

**Conclusion:** We showed that preterm birth alters cortical micro and macrostructural maturation near the time of fullterm birth. Deviations from normative development were highly variable between individual preterm infants.

## Introduction

The period encompassing the last trimester of gestation and early postnatal life is characterised by dynamic progressive and regressive cellular events that shape the developing cortex, establishing the fundamental anatomical organisation of the human brain and promoting the emergence of cognitive functioning. Rapid dendritic arborisation, synaptogenesis, ingrowth of thalamocortical afferents combined with the disappearance of radial glia and the dissolution of the subplate drive cortical growth and a significant increase in cytoarchitectural complexity (Sidman and Rakic 1973; Kostović and Jovanov-Milošević 2006). The cortex is particularly vulnerable to perturbations during this time, with deviations and/or interruptions to one or multiple of these processes predisposing to atypical development and possible long-lasting behavioural consequences (Volpe 2019).

Preterm birth, or being born before 37 weeks gestational age (GA), represents a significant risk for adverse neurodevelopment (Marlow et al. 2005). While white matter (WM) injury has been considered as the core pathology associated with preterm birth, it is now evident that altered cortical development might underlie some of the cognitive deficits in preterm birth survivors (Volpe 2005; Miller and Ferriero 2009; Rathbone et al. 2011; Fleiss et al. 2020). Magnetic Resonance Imaging (MRI) studies show that preterm infants have altered cortical growth, expansion, folding and microstructure at term-equivalent age (Ajayi-Obe et al. 2000; Ball et al. 2013, 2020; Engelhardt et al. 2015; Makropoulos et al. 2016; Bouyssi-Kobar et al. 2018; Kline et al. 2019), with a correlation between early cortical ‘dysmaturation’ and later cognitive abilities (Kline, Illapani, He, Altaye, et al. 2020). Previous work, however, has primarily focused on describing cortical development in the *preterm* brain exclusively, with little reference to normative development, or has been limited by small term-born samples. A comprehensive characterisation of the cortical sequelae of preterm birth compared to normative development is necessary to better understand the role of cortical (dys)maturation in the encephalopathy of prematurity.

In this study, we aimed to describe typical cortical micro and macrostructural maturation using *in vivo* structural and diffusion MRI in a large sample of term-born infants scanned during the neonatal period, to then test how preterm birth shapes the developing cortex. Early brain development is highly variable between individuals, and even those belonging to the same risk or clinical ‘group’ often show heterogeneous pattern of deviations from typical maturation (Dimitrova et al. 2020) and follow diverse neurocognitive trajectories (Johnson et al. 2018; Bogičević et al. 2021). Therefore, to offer a more precise understanding of the early cortical consequences of preterm birth, we not only characterise the group-average abnormality, but also capture the heterogeneity across the preterm population.

## Materials & Methods

### Participants

509 infants, of which 109 preterm born, were recruited for the developing Human Connectome Project (dHCP, approved by the National Research Ethics Committee; REC: 14/Lo/1169). Infants were scanned at term-equivalent age (TEA, 37 – 45 weeks postmenstrual age (PMA)) during natural unsedated sleep at the Evelina London Children’s Hospital. Infant preparation and imaging have been described previously in detail (Hughes et al. 2017). Exclusion criteria for the term infants are described in *Suppl. Materials.* There were no exclusion criteria for the preterm infants, except for major congenital malformations.

### MRI acquisition and preprocessing

MRI data were acquired using a 3T Philips Achieva scanner equipped with a dedicated 32-channel neonatal head coil and baby transportation system (Hughes et al. 2017). T_2_-weighted scans were acquired with TR/TE of 12s/156ms, SENSE=2.11/2.58 (both axial/sagittal) with in-plane resolution of 0.8×0.8mm (1.6mm slice thickness, 0.8mm overlap). Images were reconstructed using super-resolution and motion correction resulting in 0.5mm isotropic grid sizes (Cordero-Grande et al. 2018). Diffusion MRI (dMRI) data were acquired in a total of 300 volumes, sampling b-values of 400 s/mm^2^, 1000 s/mm^2^ and 2600 s/mm^2^ spherically uniformly distributed in 64, 88 and 128 directions, along with 20 b=0 s/mm^2^ images. Protocol parameters were multiband acceleration 4, SENSE factor 1.2, partial Fourier 0.86, acquired resolution 1.5×1.5mm (3mm slice thickness, 1.5mm overlap), TR/TE of 3800/90ms and 4 phase-encoding directions (Hutter et al. 2018).

Structural MRI data were preprocessed using the dHCP structural pipeline (Makropoulos et al. 2018). In brief, motion-corrected, reconstructed T_2_-weighted images were corrected for bias-field inhomogeneities, brain extracted and segmented into 9 tissue classes using the Draw-EM algorithm (Makropoulos et al. 2016). Following tissue segmentation, white, pial and midthickness surfaces were extracted, inflated and projected onto native left/right surfaces (Schuh et al. 2017) which were registered to a neonatal specific 40-week dHCP surface template (Bozek et al. 2018) using multimodal surface matching (MSM) (Robinson et al. 2014, 2018). For each infant, metrics of corrected cortical thickness (curvature regressed out), pial surface area (SA), curvature and sulcation were extracted (all but SA automatically generated from the dHCP pipeline).

Diffusion MRI data were preprocessed with denoising (Veraart et al. 2016), Gibbs ringing removal (Kellner et al. 2016), B_0_ field map estimation (Andersson et al. 2003) and reconstructed to 1.5mm isotropic resolution using a multishell spherical harmonics and radial decomposition signal representation with a slice-to-volume motion and distortion correction (Christiaens et al. 2021). The tensor model was fitted to the b=400 and b=1000 s/mm^2^ shells. Scalar maps of fractional anisotropy (FA) and mean diffusivity (MD), in μm^2^/ms (10^-3^ mm^2^/s), were calculated using MRtrix3 (Tournier et al. 2019). Neurite Orientation Dispersion and Density Imaging (NODDI) was fitted to all shells using the NODDI MATLAB toolbox (Zhang et al. 2012). Maps of Orientation Dispersion index (ODI) and neurite density index (or intracellular volume fraction, fICVF), were derived using the default setting in the NODDI toolbox option ‘ *invivopreterm’* with intrinsic diffusivity fixed to 1.7 μm^2^/ms and grid search starting points adjusted to better fit newborn data by lowering the range of values considered as the fraction of the intra-neurite space from 0-1 to 0-0.3 (Kunz et al. 2014).

Infants’ dMRI-derived maps were affine registered to native T_2_-weighted images via the mean b1000 shell using *epi_reg,* implemented in FSL (Jenkinson et al. 2002). When projecting diffusion metrics to the cortical surface, we applied partial volume correction by estimating partial voluming on a voxel level using Toblerone (Kirk et al. 2020) *(Suppl. Materials & Suppl. Fig. 1).* Following partial volume estimation and correction, diffusion maps were projected to native surface space using ribbon constrained mapping and resampled to the 40-week dHCP neonatal surface template using adaptive barycentric interpolation (implemented in Connectome Workbench, https://www.humanconnectome.org/software/connectome-workbench).

We used a two-stage quality control (in volume and surface space) to ensure no data affected by (i) motion, or with (ii) artefacts, (iii) poor dMRI to T_2_-weighted registration, (iv) unsatisfactory surface projection or registration to surface template were included *(Suppl. Material).* The final sample comprised 259 term-born and 76 preterm infants *(Suppl. Table 1).* Cortical regions were defined using approximately equal-sized cortical parcellations with Voronoi decomposition (n=143 per hemisphere). For all metrics with the exception of SA, where the sum was used, we calculated the mean value per parcel for every infant to be used for subsequent analyses.

### Typical cortical development in term-born infants

#### Age-associated change in cortical morphology

First, we described age-associated changes in cortical morphology in the full 259 term-born sample. This was performed with Permutation Analysis of Linear Models (PALM) implemented in FSL using 10000 permutations and a family-wise error (fwe) correction across modalities (8 metrics) and contrasts (positive/negative association with PMA at scan), including sex in the model (Winkler et al. 2014; Alberton et al. 2020). Pearson’s coefficient was reported for parcels showing significant correlation at p_mcfwe_ < 0.05 (***m***odality ***c***ontrast***fwe*** corrected).

#### Predicting age at scan using cortical features

In addition, to predict PMA at scan, we used Random Forest (RF) regression in *Scikit Learn* (Pedregosa et al. 2011). The 259 term-born infants were split into training (75%) and hold-out (25%) samples *(Suppl. Fig. 2).* Hyperparameters were tuned on the training set using Bayesian Optimisation search *(Scikit Learn Optimize)* under 5-fold cross validation and mean absolute error (MAE) as loss function. Optimal parameters were max_depth = 35, max_features = ‘auto’, min_sample_leafs = 2, n_estimators = 1000. The RF predictions were also error-in-variables bias corrected using linear regression (Smith et al. 2019). Model performance was evaluated by estimating the MAE, mean squared error (MSE) and Spearman correlation (ρ) between true and predicted PMA at scan in the hold-out sample using the RF and error-in-variables correction models estimated on the training data. We examined the top 10% of the features using the default *Scikit Learn feature_importances_.*

### Cortical development in the preterm brain at term-equivalent age

#### Group-level differences between preterm and term infants

We tested for group-average differences between term (full 259 term-born sample) and preterm infants (n=76). Aggregate values (mean and sum for SA) were calculated for every parcel and infant. Analyses were performed in PALM, correcting across contrasts (preterm < term; preterm > term) and modalities (8 metrics). We report the t-statistic for parcels indicating a significant group mean difference at p_mcfwe_ < 0.05.

#### Mapping cortical variability in the preterm brain with Gaussian process regression

To capture the heterogeneity of cerebral development in the preterm brain, we used multioutput Gaussian process regression (GPR), a Bayesian non-parametric regression, implemented in GPy (https://sheffieldml.github.io/GPy/). GPR simultaneously provides point estimates and measures of predictive confidence for every observation representing the distance of each individual from the normative mean at that point on the ‘curve’ accounting for modelled covariates. Analogous to the widely employed paediatric height and weight growth charts, this technique allows the local imaging features of individual infants to be referred to typical variation while simultaneously accounting for variables such as age and sex (Marquand et al. 2016, 2019; Wolfers et al. 2018; O’Muircheartaigh et al. 2020).

We first trained a GPR model to describe normative cortical development in the training term-born sample (75% of the term sample: 196 infants; *Suppl. Fig. 2)* using PMA at scan and sex as predictors and FA, MD, ODI, fICVF, cortical thickness, SA, curvature and sulcation as outputs. The relationship between cortical outputs and model predictors was estimated with a sum of radial basis function, linear and white noise covariance kernels. Model hyperparameters were optimised using log marginal likelihood. Prediction performance was evaluated using the MAE between predicted and observed values derived from the held-out term-born sample (25% of the term sample: 63 term infants). To assess the effects of preterm birth, we applied the model to 76 preterm infants scanned at TEA. A Z-score was derived for every infant in the preterm and hold-out term-born samples by estimating the difference between model prediction and observed value normalised by the model uncertainty (the square root of the predicted variance). To quantify extreme deviations, prior to analyses, we chose a threshold of |Z|>3.1 (corresponding to p<0.001) (Dimitrova et al. 2020).

#### Spatial overlap maps

The spatial prevalence of extreme negative/positive deviations across the preterm sample was examined by calculating a parcel-based percentage map of extreme deviations (number of infants with |*Z*| > 3.1 divided by the total number of infants), individually for every cortical feature.

#### Atypicality index

For every cortical feature and infant, we calculated a positive and a negative whole-cortex atypicality index. These indices capture the extreme positive/negative deviations for every infant and represent the ‘overall burden’ as a percentage of cortical parcels with extreme deviations relative to the total number of parcels.

#### Deviations from normative development, age at birth and behaviour at 18 months

To test if age at birth influences cortical development, we examined the association between GA at birth and GPR derived Z-scores; and between GA at birth and the whole-cortex atypicality indices (in preterm infants; in hold-out term infants and in the combined preterm and hold-out term-born infants). To assess if deviations from normative cortical development relate to later behaviour, we looked at the correlation between Bayley III Scales of Infant and Toddler Development (BSID-III) (Bayley 2006) composite motor, cognitive and language scores, collected at 18 months corrected age, and Z-scores and atypicality indices. PALM was used (as described above) for parcel-wise Z-score data; Kendall *τ* (for correlation) and Mann-Whitney *U* (for group differences) for the atypicality indices. Analyses were corrected for multiple comparison correction (FDR).

### Data availability

The MRI data for the dHCP are available at http://www.developingconnectome.org/. dHCP surface template is available at https://brain-development.org/brain-atlases/atlases-from-the-dhcp-project/cortical-surface-atlas-bozek/. Code for MSM surface registration can be found at https://github.com/ecr05/MSM HOCR, partial volume correction code at https://git.fmrib.ox.ac.uk/seanf/dhcp-neonatal-fmri-pipeline/-/blob/90f2f7317fff0cd809d0b1b4a400d970c1ea115c/doc/surface.md. Python code to perform the main analyses in this work will be available from https://github.com/ralidimitrova.

## Results

Term and preterm infants did not differ in PMA at scan (p=0.90), nor in sex distribution (p=0.89) (Table 1). Distribution of GA at birth and PMA at scan are shown in *Suppl. Fig. 3*. While punctate white matter lessions (PWMLs) were more frequently seen in the preterm sample (p<0.05), no infants had severe WM injury such as periventricular leuomalacia and only one infant had more than 20 PWMLs. One preterm infant had cystic lesions in both thalami extending to internal capsule and left putamen, one had cerebellar hemispheric parenchymal loss accompanied by brainstem atrophy, and one had middle cerebral artery infarct (the last resulting in gross cortical injury, excluded from all group-level analyses). No group differences were observed at the 18 month follow-up (cognitive p=0.99; motor p=0.19; language p=0.36). All, but two (one in language, one in cognitive outcome), preterm infants scored within 2 standard deviations from the population mean.

**Table 1.**
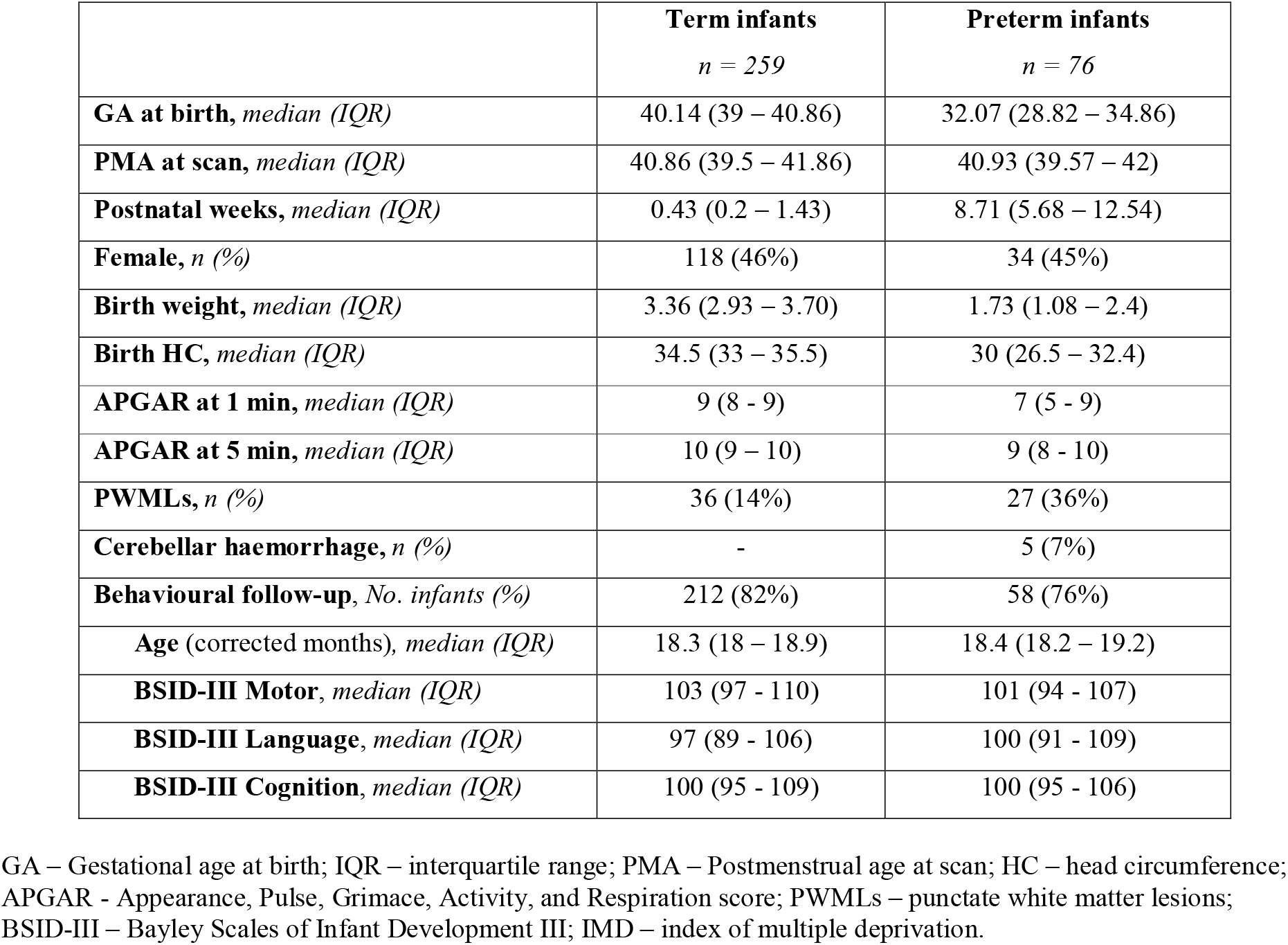
Characteristics of the study sample.

### Rapid regionally specific micro and macrostructural development in the term-born neonatal cortex

We observed a rapid change in cortical micro- and macrostructural composition during the neonatal period (Fig. 1). Overall, there was a reduction in FA in the parietal, occipital and temporal lobes and in MD in the central sulci, the anterior cingulate, the insula and in some parcels located in the frontal cortex. There was an increase in ODI across the brain, steepest in the parietal and temporal lobes, but no change in the somatosensory cortex. We also observed an increase in fICVF mainly in the insula but also in some regions of the frontal lobe. Cortical thickness increased in the somatosensory, occipital, temporal, insular and portions of the parietal and frontal regions. SA increased across the whole cortex, showing the most dramatic age-associated change compared to all other metrics. While curvature remained stable over the studied period, cortical sulcation increased in the cingulate cortex.

**Figure 1.**
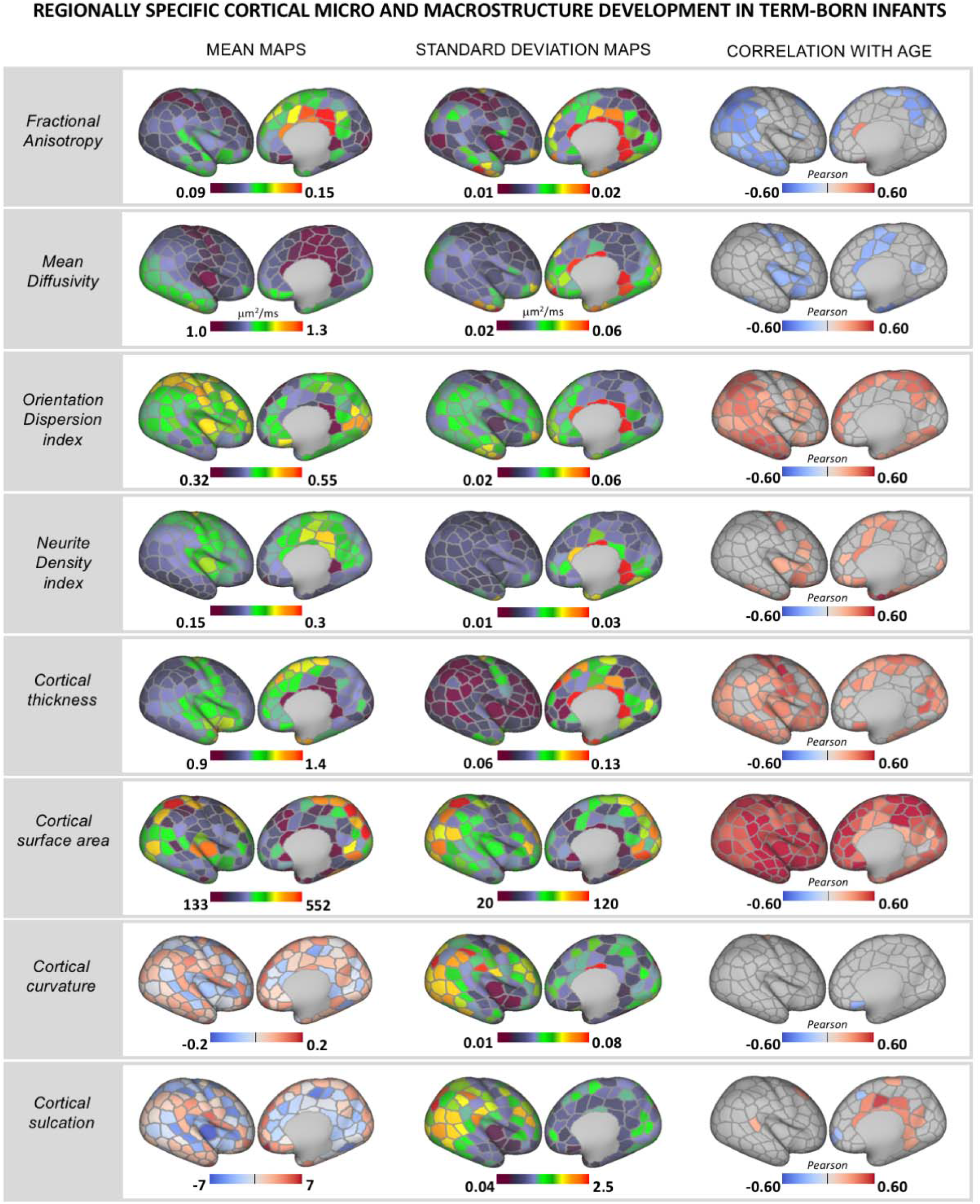
Rapid regionally specific cortical micro- and macrostructure development in term-born infants during the neonatal period. Mean and standard deviation surface maps are shown for all metrics together with Pearson’s correlation coefficient for all parcels showing significant (p_mcfwe_< 0.05) positive (red) or negative (blue) association with age (PMA at scan). Right hemisphere depicted.

RF regression predicted PMA at scan in the held-out term-born sample with MAE = 0.60, MSE = 0.56 (weeks) with correlation between observed and predicted age of ρ = 0.85, R^2^ = 0.74 *(Suppl. Fig. 4).* SA was the ‘best’ predictor of PMA at scan, with 47% (108/229 parcels) of the 10% most important features. ODI followed with 37 parcels (16%), FA with 14 (6%), cortical thickness with 29 (13%), fICVF with 21 (9%), MD with 11 (5%), sulcation with 6 (3%) and curvature with 3 parcels (1%) *(Suppl. Fig. 5).*

### Cortical consequences of preterm birth at term-equivalent age

#### Group-level differences between term and preterm infants

At a group level, preterm infants showed higher MD and lower fICVF across a large proportion of the posterior cortex, including parcels in the parietal, occipital and temporal lobe (Fig. 2A). In the insula, preterm infants had higher FA and lower ODI compared to term-born peers. We also observed greater cortical thickness in frontal, insular and anterior parietal cortices in preterm compared to term-born infants. Apart from a few parcels, there were no group differences in SA, curvature, or sulcation.

**Figure 2.**
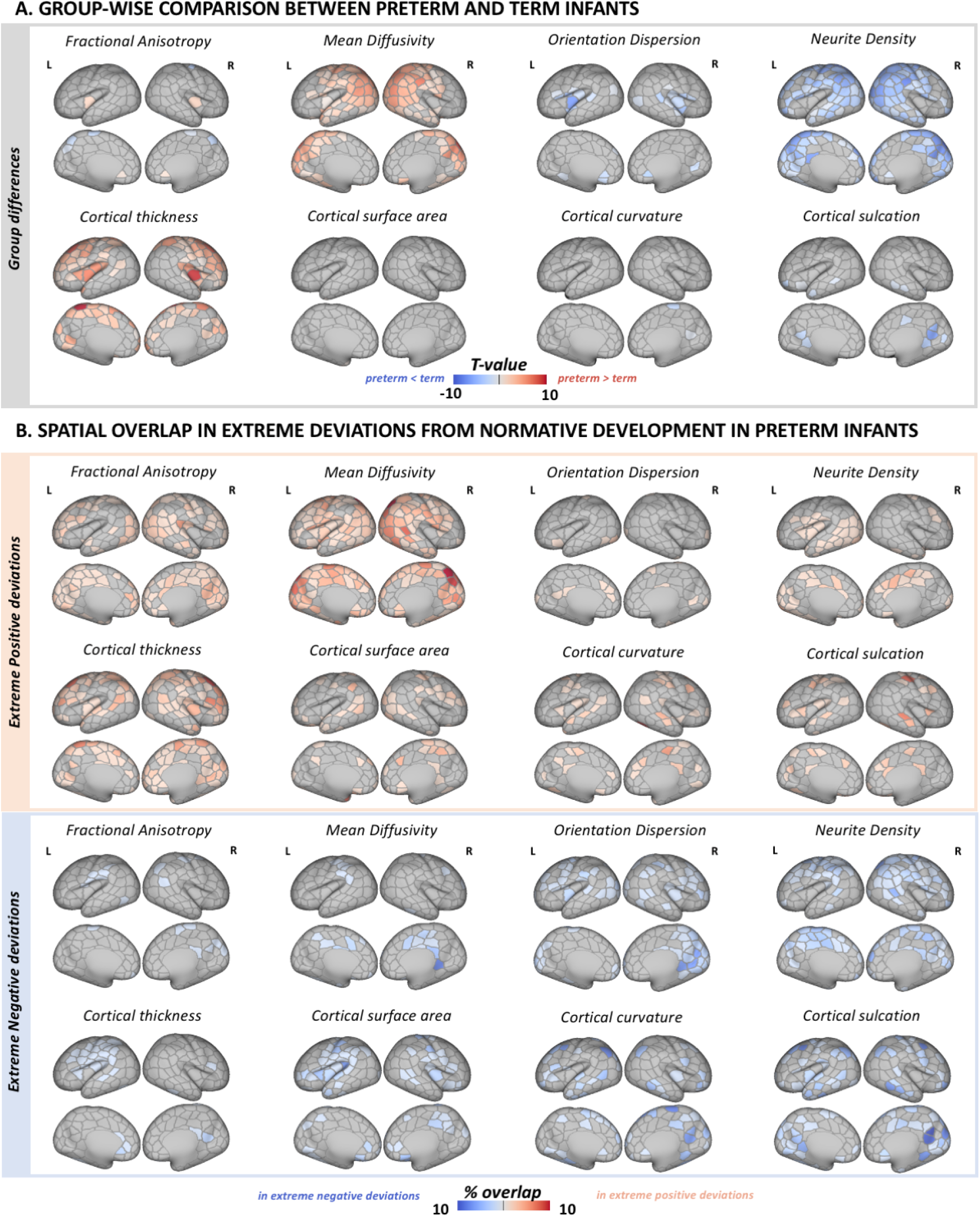
Effect of preterm birth on the developing cortex. (A) Group-level differences in cortical macro- and microstructure between term and preterm infants. Depicted are only parcels that show significant group-wise differences at p_mcfwe_ < 0.05. (B) Spatial overlap in extreme positive (red) and negative (blue) deviations from normative development in preterm infants. The overlap maps show the proportion of infants with extreme deviations (Z > |3.1|) from normative development for every parcel.

#### Heterogeneity in individual extreme deviations from normative development in the preterm sample

We observed widespread extreme positive deviations in MD and thickness, and extreme negative deviations in fICVF in preterm infants (Fig. 2B). While there was a visible overlap in parcels showing group differences in these metrics and regions with individual extreme deviations, in some cortical areas we detected only group-average alterations but no individual extreme deviations in preterm infants. And vice versa, in other parcels we observed extreme deviations on an individual infant level but not in the group-level analyses. For example, very few parcels showed group differences in FA, ODI and SA, but the spatial overlap maps were indicative of widespread individual deviations in extreme positive FA, negative ODI and positive/negative SA deviations. Overall, there were very few parcels across all metrics with more than 8% of the infants showing extreme deviations highlighting the high variability in cortical development associated with preterm birth. Term-born infants showed fewer and less overlap in extreme deviations compared to preterm infants *(Suppl. Fig. 6).* Model MAE maps are shown in *Suppl. Fig. 7.*

#### Parcel-level deviations from normative development and age at birth

Lower GA at birth related to increased MD and decreased fICVF in the parietal cortex (more prominent in the left hemisphere; Fig. 3.), and increased FA (bilaterally) and lower ODI (left hemisphere) in the insula. We observed a negative correlation between GA at birth and cortical thickness in the bilateral frontal and insular cortex. Very few parcels showed an association with GA at birth in the preterm or term samples alone (*Suppl. Fig. 8*). We observed no association between Z-scores and BSID-III neurodevelopmental scores at 18 months.

**Figure 3.**
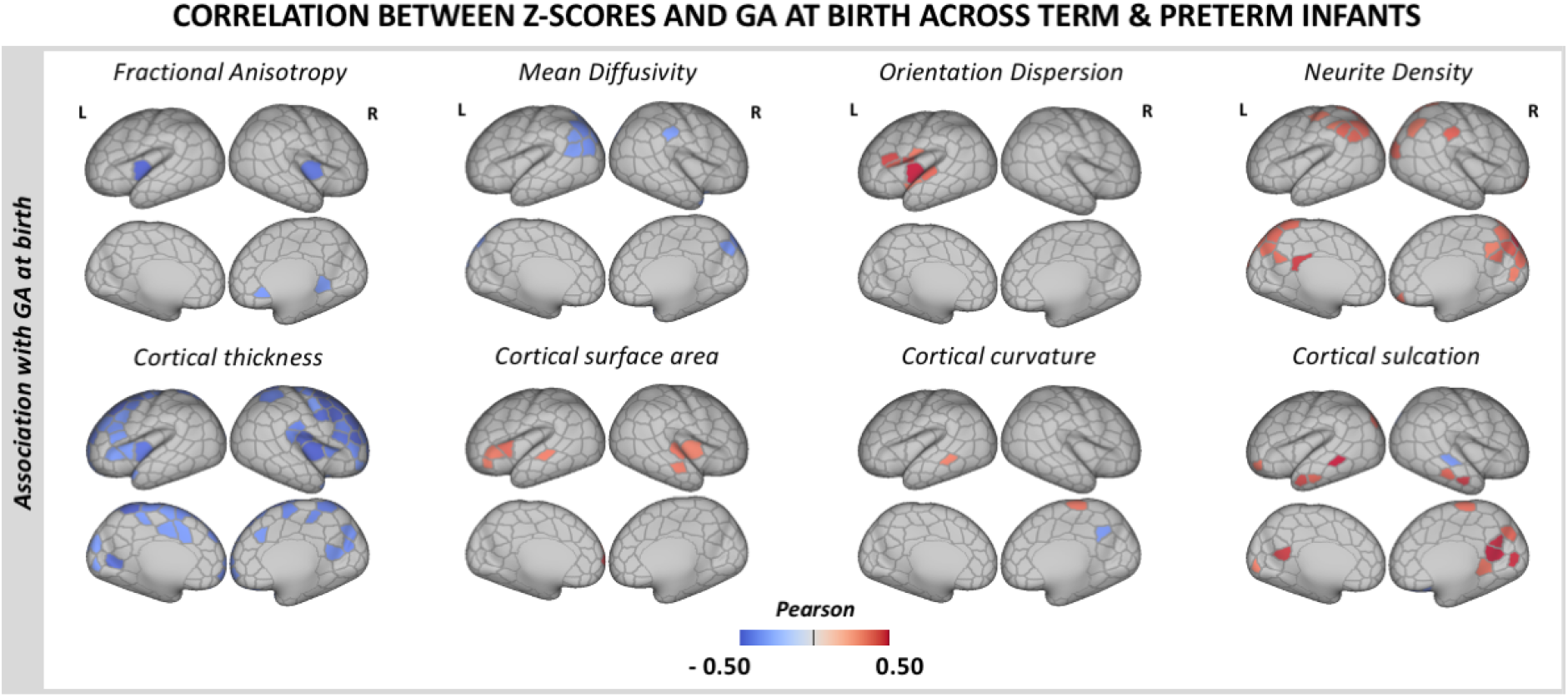
Association between deviations from normative cortical development (Z-scores) and GA at birth. Pearson’s correlation coefficient is shown only for parcels showing significant correlation with GA at birth at p_mcfce_< 0.05 in the combined term-born hold-out and preterm samples.

#### Whole cortex atypicality index

As a group, preterm infants had higher atypicality indices for all metrics except FA (Fig. 4). On average, the preterm brain had higher proportion of extreme negative deviations in ODI, fICVF, curvature and sulcation, as well as higher proportion of extreme positive deviations in MD, thickness and sulcation (all at p_FDR_<0.05). Term-born infants did not have higher atypicality index in any of the cortical features we studied.

**Figure 4.**
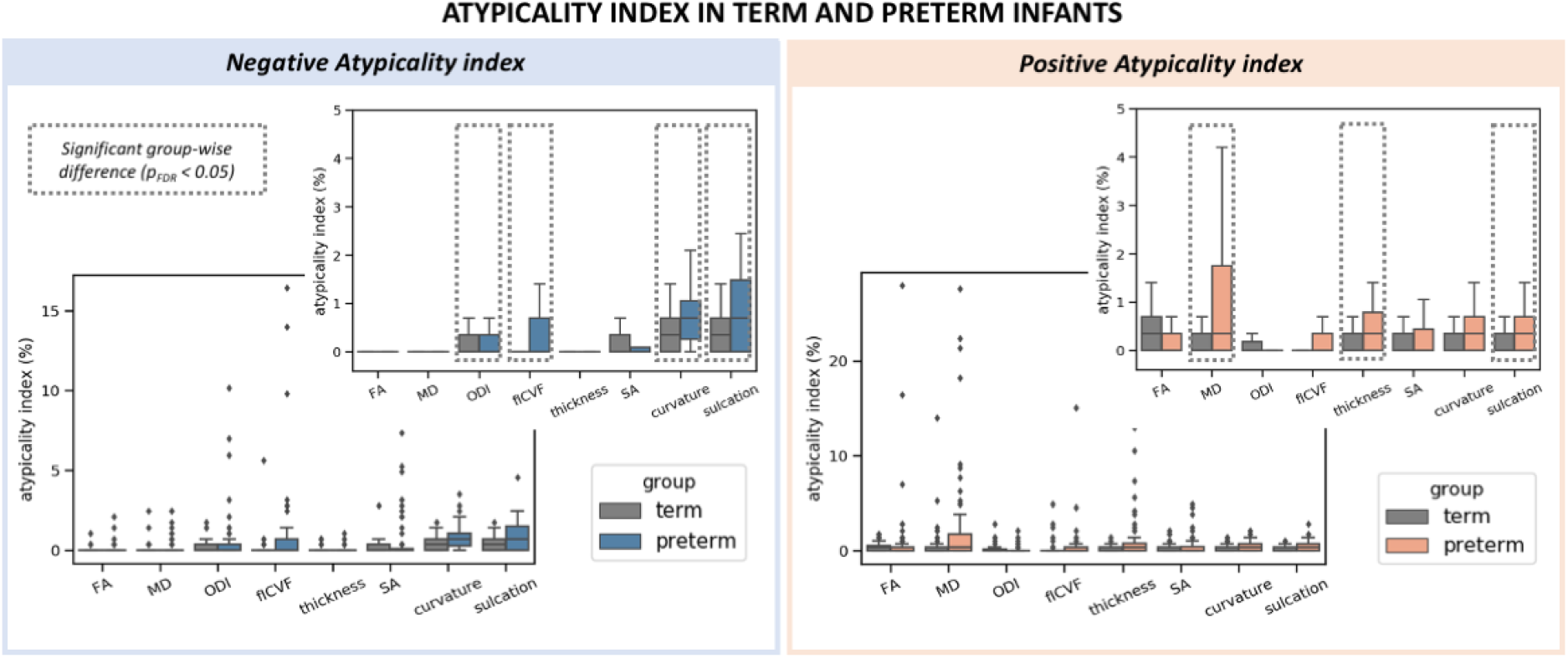
Distribution of the atypicality indices in term and preterm infants. The preterm brain showed higher burden of both negative (left) and positive (right) extreme deviations. Boxplots are shown with and without outliers, the latter to highlight the distribution of the indices in the two groups. Indices where a group-level difference was found are highlighted.

Lower GA at birth was related to higher proportion of extreme negative deviations in ODI (= −0.15), fICVF (= −0.26), curvature (= −0.24) and sulcation (= −0.23), all at p_FDR_<0.05). Being born earlier was also associated with higher proportion of extreme positive deviations in MD (= −0.24), thickness (= −0.20) and sulcation (= −0.19), all at p_FDR_<0.05).

After multiple comparison correction, we observed no significant association between the atypicality indices and BSID-III scores at 18 months in the preterm sample alone, nor in the combined term-born hold-out and preterm samples.

## Discussion

During early brain development, the cortex undergoes rapid changes that establish the fundamental anatomical organisation of the brain. Within the period between 37 and 45 weeks PMA, we observed regionally specific micro and macrostructural cortical maturation in a large sample of term-born infants. Abrupt exposure to extra-uterine environment through preterm birth resulted in atypical cortical microstructure and growth at TEA, with marked variability in individual deviations from normal development. Deviations in regional cortical maturation were associated with GA at birth but not with neurodevelopment at 18 months.

### Dynamic and spatially heterogeneous maturation of the term-born neonatal cortex

By the end of full-term gestation, neuronal migration is mostly completed, thalamocortical afferents have reached their cortical targets and the basic columnar organisation is largely established (Sidman and Rakic 1973; Kostović et al. 2002; Kostović and Jovanov-Milošević 2006; Paredes et al. 2016). After birth, the dramatic increase in dendritic arborisation and synapse production underlies the development of more complex cytoarchitecture and contributes to rapid cortical growth (Marin-Padilla 1967; Huttenlocher and Dabholkar 1997; Volpe 2019). During this period, we observed increasingly more complex microstructural ‘geometry’ in the neonatal term-born brain, captured by changes in anisotropy (FA↓) and neurite orientation dispersion index (ODI↑), accompanied by reduction of tissue water content (MD↓) and increase in neurite density index (fICVF↑). A fall in anisotropy (FA↓) and increase in orientation dispersion index (ODI↑) were present in parietal, occipital and temporal regions, with increasing ODI also evident in the frontal lobe. Neurite density index increased (fICVF↑) and tissue water content decreased (MD↓) in somatosensory and insular cortices. These changes suggest regionally and temporally asynchronous cortical maturation after birth (Yu et al. 2016; Batalle et al. 2019; Ouyang et al. 2019; Fenchel et al. 2020). The newborn brain is exposed to an environment rich in sensory stimuli that promotes synaptogenesis and dendritic arborisation postnatally, especially in sensory areas (Huttenlocher 1990; Huttenlocher and Dabholkar 1997), which might account for the rapid age-associated change we observed in these regions.

We described different spatiotemporal patterns of SA and cortical thickness growth. While surface expansion was very rapid and relatively uniform across the developing cortex, cortical thickness increased in the somatosensory, temporal, occipital, insular and parts of the frontal lobe. We observed a steeper age-associated change in SA compared to cortical thickness (Lyall et al. 2015; Jha et al. 2019) and proportionally, SA features were the ‘best’ predictors of age at scan in our RF model. Very little change in curvature and sulcation was observed, suggesting that folding patterns are largely established by full-term gestation (Chi et al. 1977; Hill et al. 2010; Meng et al. 2014; Li et al. 2015; Batalle et al. 2019; Fenchel et al. 2020). These findings further support the complex asynchronous pattern of cortical growth in both primary and association regions during early postnatal development (Jha et al. 2019).

### Preterm birth alters the developing cortex at term-equivalent age

The dynamic nature and complexity of the cellular events that take place in the last trimester makes the developing cortex particularly vulnerable to perturbations (Volpe 2019). A number of studies have characterised the developmental trajectory of cortical microstructure in the *preterm* brain, but only a few offer a direct reference against a large normative dataset (McKinstry et al. 2002; Ball et al. 2013, 2020; Eaton-Rosen et al. 2015, 2017; Smyser et al. 2016; Yu et al. 2016; Batalle et al. 2019; Ouyang et al. 2019). Our results indicate that the *preterm* cortex is more ‘immature’ at the time of full-term birth. Specifically, we show that *on average* preterm infants have increased tissue water content (MD↑) and reduced neurite density index (fICVF↓) in the occipital, parietal, temporal and the somatosensory cortex compared to term-born peers. Group differences in cortical microstructural ‘geometry’ were evident in few parcels, mainly in the insular region, suggesting reduced neurite orientation complexity (FA↑ and ODI↓). GA at birth was negatively associated with MD and positively with fICVF in the parietal cortex, while in the insular cortex increased GA related to lower FA and higher ODI. The exact cellular mechanisms underlying this altered microstructure development are unknown. Dysmaturation of the neurons in the subplate, that offer initial targets and guidance of thalamocortical afferents before they reach the cortical plate may have a contributing role (Volpe 1996, 2019). Disrupted dendrite and spine formation, or a general reduced morphological complexity of cortical neurons, is another possible cause (Dean et al. 2013).

On average, preterm infants had thicker cortices in the somatosensory, frontal and insular regions. We observed a negative association between cortical thickness and GA at birth in most of these regions, an association that seems to persist into infancy (Jha et al. 2019). Greater cortical thickness in preterm individuals has been observed in childhood and adolescence, arguing atypical trajectory of cortical thinning as differences with term-born controls substantially decrease in early adulthood (Mürner-Lavanchy et al. 2014; Nam et al. 2015). Although the exact mechanisms behind this atypical cortical growth in the preterm brain are largely unknown, they likely include altered neuronal differentiation or modification/pruning of early overproduced neurons and their processes in the cortical plate. We did not observe group-level differences in SA as reported by others (Lax et al. 2013; Engelhardt et al. 2015; Zhang et al. 2015), nor did we see a strong correlation between SA and GA at birth (Jha et al. 2019), apart from the insular cortex. SA and cortical thickness are genetically independent, determined by distinct cellular processes (Rakic 1995; Panizzon et al. 2009) and follow different and regionally heterogeneous developmental trajectories (Lyall et al. 2015). While the number of cortical minicolumns established during embryonic development is believed to determine SA, the number, size and density of neurons and their processes within each minicolumn produced during the foetal and perinatal period, are thought to impact cortical thickness (Rakic 1995, 2009). The timing of the abrupt exposure to extrauterine environment might explain why cortical thickness appears more affected than SA in preterm infants.

### Group-average atypicality and individual variability in the preterm cortex at term-equivalent age

While we show a good agreement between group-wise analyses and spatial overlap maps in extreme deviations from the normative development in MD, fICVF and thickness, high variability across individual preterm infants was evident in all metrics (Fig. 2). Very few cortical parcels had more than 8% of the preterm group with extreme deviations in a given metric. These findings are consistent with our previous work describing heterogeneous volumetric and wholebrain microstructure development at TEA following preterm birth (Dimitrova et al. 2020, 2021). This highlights that group level understanding of the preterm brain disguises a large degree of individual variability, that may be important to single infants. Preterm infants were more likely to show higher proportion of parcels with extreme deviations from normative development (atypicality indices) compared to term-born infants (Fig. 4). Being born earlier was associated with higher proportion of parcels with extremely low (Z < −3.1) ODI, fICVF, curvature and sulcation, and with more parcels with extreme high (Z > 3.1) MD, thickness and sulcation. This argues that when reaching the time of full-term birth, the cortex is more immature in those born at lower GA. While we did not detect mean differences in curvature and sulcation between the two groups, preterm infants were more likely to show higher loading of extreme deviations from the normative curvature and sulcation compared to term infants. Atypical folding pattern has been previously reported in preterm infants (Engelhardt et al. 2015; Zhang et al. 2015) and our study highlights the high variability associated with it.

### Cortical deviations were not associated with later neurodevelopment

Preterm birth, especially before 25 weeks GA, has been linked to poor outcome in childhood (Wood et al. 2000; Marlow et al. 2005), with a strong correlation between degree of prematurity and later cognitive functioning (Bhutta et al. 2014). Several cortical features including curvature, SA and cortical microstructure have shown promise as biomarkers predictive of later neurodevelopment in preterm infants (Kapellou et al. 2006; Rathbone et al. 2011; Ball et al. 2013; Kline, Illapani, He, Altaye, et al. 2020). While we report atypical cortical development in the preterm brain at TEA, we did not observe an association between individual deviations from normative development and 18-months follow-up, or a strong correlation between cortical features and GA at birth in preterm infants. Compared to previous datasets primarily comprising extremely and/or very preterm infants including infants with poor outcome (Ball et al. 2013; Kline, Illapani, He, Altaye, et al. 2020; Kline, Illapani, He, and Parikh 2020), our sample also included mid and late preterm infants and all infants, but two, showed normative outcome at 18 months (Table 1). In a previous study of this cohort, we found an association between overall atypicality index of whole-brain microstructure, mainly in the developing WM, and outcome (Dimitrova et al. 2020). In the preterm brain at TEA, even in the lack of overt injury, WM dysmaturation may have a stronger impact on later behaviour compared to deviations in cortical development (Woodward et al. 2006).

### Limitations

Direct correlation between imaging and histological examinations of human brain development are sparse (Trivedi et al. 2009; Dean et al. 2013; Huang et al. 2013), and caution is crucial when contributing changes in any diffusion metric to a specific biological process(es) (Alexander et al. 2007; Kroenke 2018; Kamiya et al. 2020). Here we combined DTI and NODDI, the two most widely applied dMRI models in the study of *cortical* microstructure. DTI is highly sensitive to underlying microstructure, however, inherently lacks specificity. In particular, FA is modulated by multiple factors including dendritic/axonal density, number, orientation dispersion and partial volume contamination, creating a considerable challenge in attributing changes in FA to (a) specific aspect(s) of the underlying architecture (Jones et al. 2013). While NODDI promises higher biological specificity, the model carries several limitations including the absence of any direct intrinsic diffusivity estimation. The decision to fix the intra and extracellular parallel diffusivity to the same predetermined value might represent a major source of systematic oversimplification and lead to a potential bias in parameter estimation reducing their promised specificity (Jelescu et al. 2016; Jelescu and Budde 2017). While recent work has shown significant promise (Guerrero et al. 2019), the optimal values for model parameters used in the study of the *developing* cortex are yet to be established. We chose to use the default *‘invivopreterm’* setting in NODDI to be consistent with previous studies (Batalle et al. 2019; Ball et al. 2020; Fenchel et al. 2020). However, more work is essential to validate the estimation of the model parameters and their biological accuracy.

During the neonatal period, brain morphology is rapidly changing, the cortex is only approximately 1.1mm thick and tissue contrast changes dramatically with maturation (Makropoulos et al. 2018; O’Muircheartaigh et al. 2020). This makes image registration and segmentation of the developing brain particularly challenging. Compared to the traditional volume-based registration, the surface-based method we used is driven by geometric features of cortical shape and thus offers superior inter-subject cortical alignment (Robinson et al. 2014, 2018; Bozek et al. 2018) and lessen CSF contamination (Glasser et al. 2016). To further ensure partial volume effects are minimised, we applied surface-based partial volume correction of the dMRI data. We also used a conservative two-stage QC to ensure only data of good quality and successful preprocessing were used, and thus we are confident that we report biological effects.

## Conclusion

We described rapid and spatially varying cortical micro and macrostructure maturation in the term-born neonatal brain. We also showed that abrupt interruption to gestation through preterm birth alters cortical development near the time of full-term birth and deviations from normative development are variable across individual preterm infants.

## Acknowledgements

The authors would like to thank all the families who dedicated their time to take part in the study. The authors declare no conflict of interest.

