## Supplementary material for "Preterm birth alters the development of cortical microstructure and morphology at term-equivalent age": Suppl

*Exclusion criteria for term infants*:

Exclusion criteria for the term-born sample included admission to neonatal intensive care unit or significant intracranial abnormality detected on neonatal MRI scan (including acute infarction and parenchymal haemorrhage), but not punctate white matter lesions (PWMLs), small subependymal cysts/haemorrhages in the caudothalamic notch, mildly prominent ventricles or widening of the extra-axial CSF (within normative variation).

*Partial volume correction*

Despite achieving 1.5 mm isotropic voxel resolution in the dHCP, the dMRI data voxel resolution remains larger than the estimated ~1.1mm neonatal cortical thickness (Makropoulos et al. 2018). We applied partial volume correction (PVC) by estimating partial voluming on a voxel level using Toblerone (Kirk et al. 2020). This resulted in 3 tissue probability maps (GM, WM, CSF). Voxels with cortical probability < 5% were masked out from the projection to the surface. *Supplementary Figure 1* shows the difference between MD surface with and without PVC for a single infant (upper panel) and mean group average for the term-born sample (lower panel).

*Quality control for dMRI and sMRI data*:

The initial sample consisted of 400 term and 109 preterm infants scanned at TEA with acquired dMRI and structural MRI (sMRI) data. We used a two-stage quality control (QC). Exclusion of subjects is broken down in *Supplementary Table 1.* Initial QC was performed in volume space and examined the success of the sMRI and dMRI preprocessing pipelines. QC of the sMRI data included visual examination of motion corrected T_2_-weighted images following a scoring method described in (Makropoulos et al. 2018). For dMRI data, nominal metrics of the total amount of motion were estimated from the output translation and rotation trajectories of every subject. These translation and rotation metrics were respectively defined as the root sum of squares of the forward difference of the translation parameters (in mm) and the rotation parameters represented in Lie algebra (Claraco, 2018). When combined, they relate to the total length of the subject’s motion trajectory during the entire dMRI sequence. A summary motion QC metric was quantified for every infant following the procedure described in (Christiaens et al. 2021), and data of insufficient quality were excluded (QC score > 3.5). Further visual assessment confirmed the success of preprocessing, alignment to T_2_-weighted native space and ensured that all brain data were within the field of view (i.e., dMRI data where part of the cortex was cropped due to the infant moving out of the field of view were excluded). The second QC was performed in surface space and involved visual examination of individual surfaces ensuring successful projection, no gross misalignment to the 40-week dHCP surface template or cropped cortex not noted in volume space. Seventeen term infants were excluded from the term sample due to incidental findings. The final sample consisted of 259 term and 76 preterm infants.

**Supplementary Figures**

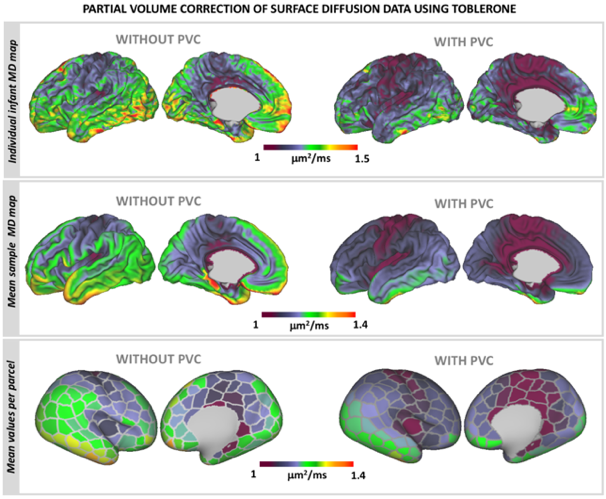

***Supplementary Figure 1.*** Partial volume correction (PVC). Individual infant Mean Diffusivity surface map without and with PVC (upper panel). Average Mean Diffusivity (MD) map without and with PVC for the term-born sample (lower panel).

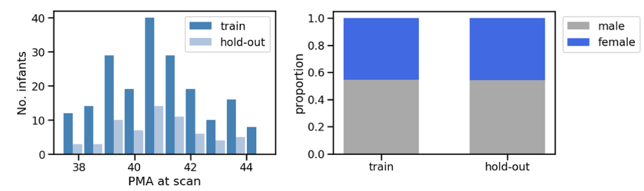

***Supplementary Figure 2.*** Distribution of PMA at scan and sex proportion between train and hold-out term-born sample split used in the RF and GPR modelling.

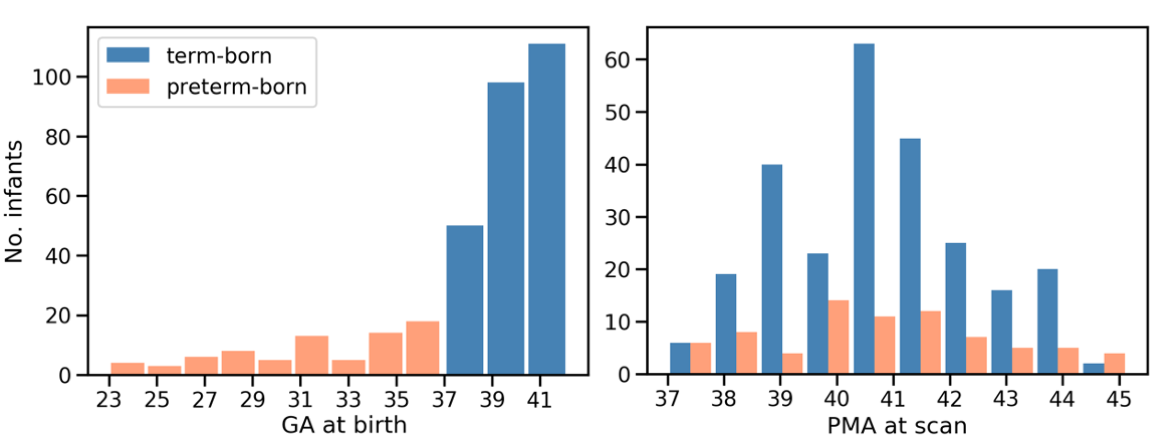

***Supplementary Figure 3.*** Distribution of GA at birth and PMA at scan in the term and preterm samples.

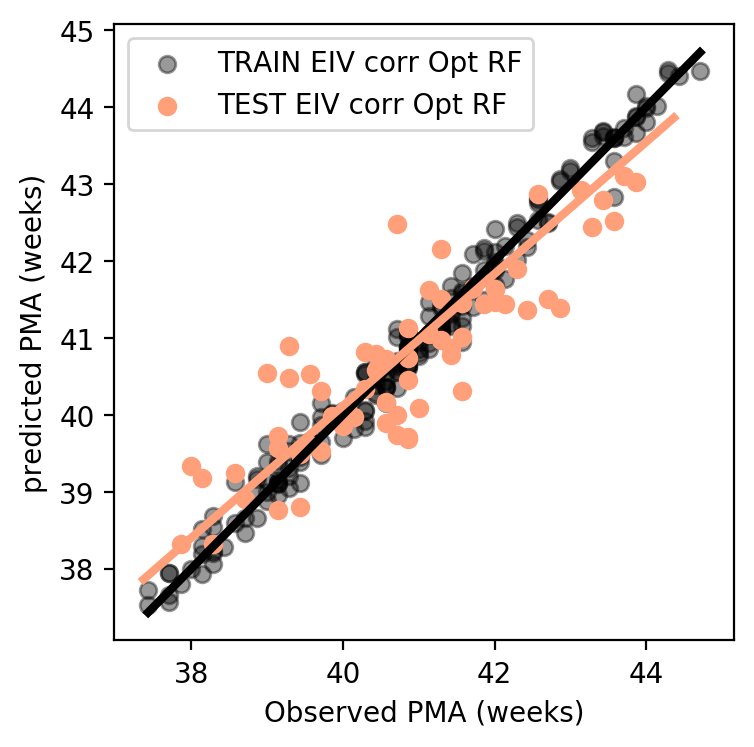

***Supplementary Figure 4.*** Predicted versus observed age at scan (PMA) for the train and hold-out samples with best line of fit for the error-in-variables (EIV) hyperparameter optimised RF model.

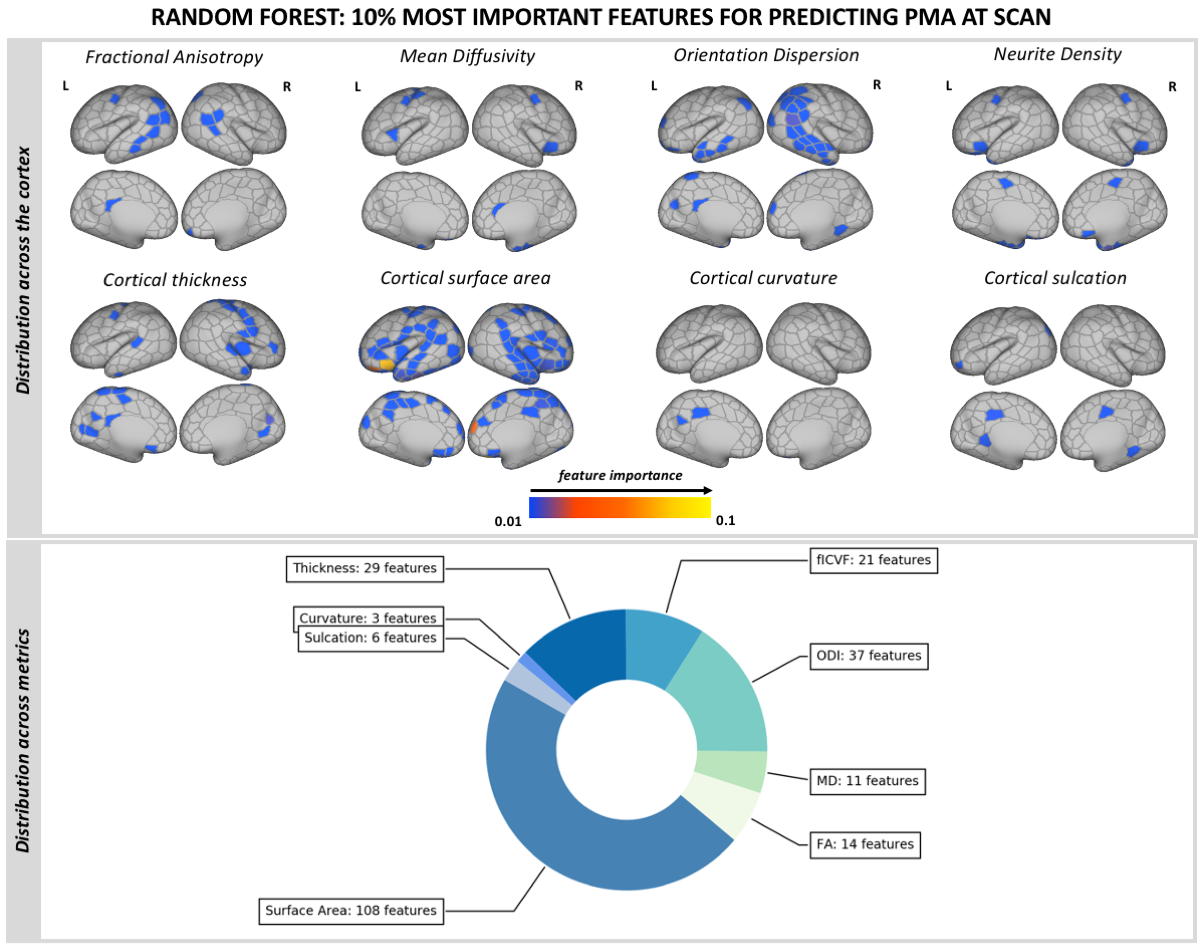

***Supplementary Figure 5.*** Predicting PMA at scan using cortical surface features in term-born infants using RF regression. The 10% most important features (229) are shown.

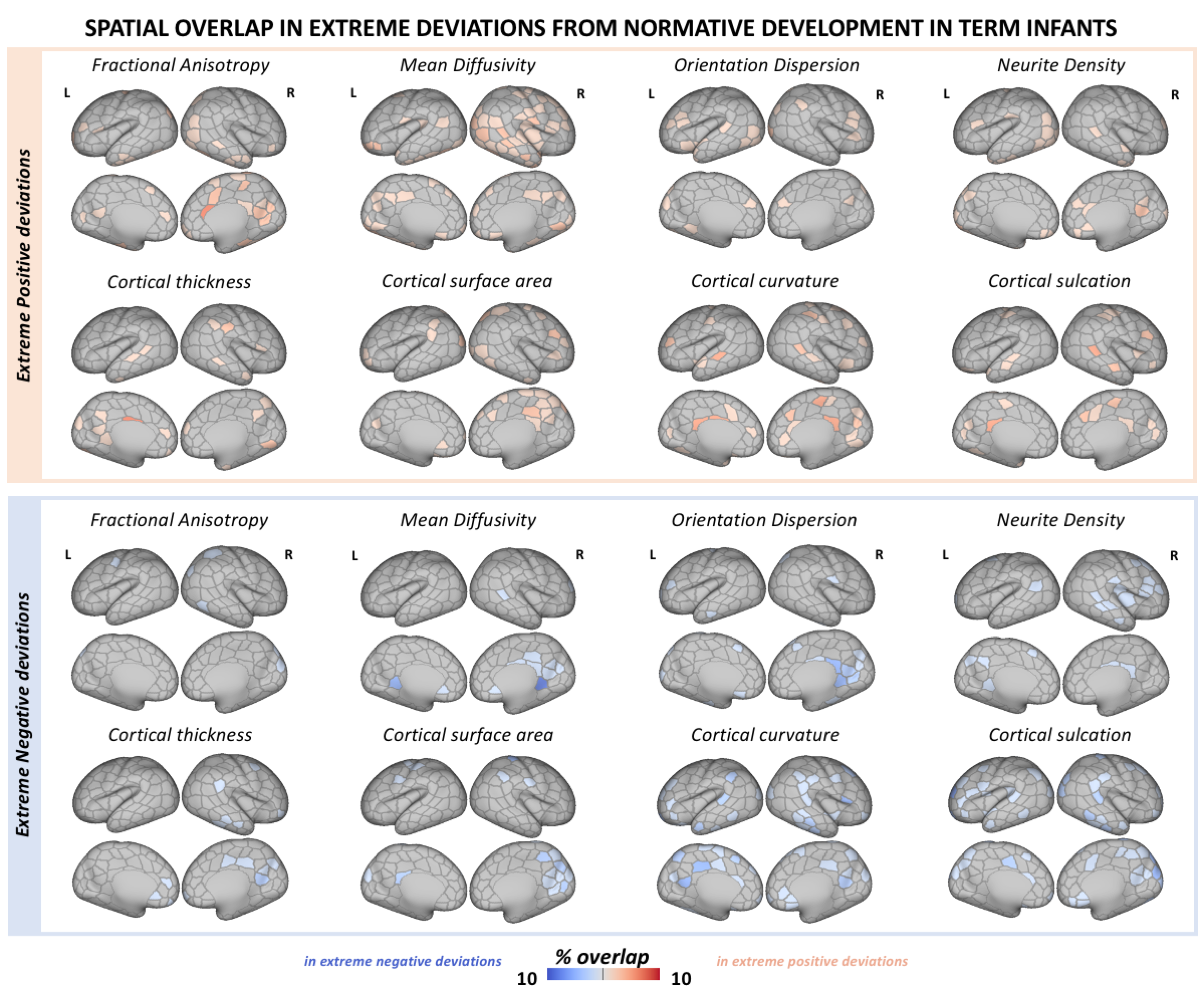

***Supplementary Figure 6.*** Spatial overlap in extreme positive/negative deviations from normative development in the hold-out test term-born infants. The overlap maps show the proportion of infants with extreme deviations (Z > |3.1|) from normative development for every parcel and cortical feature.

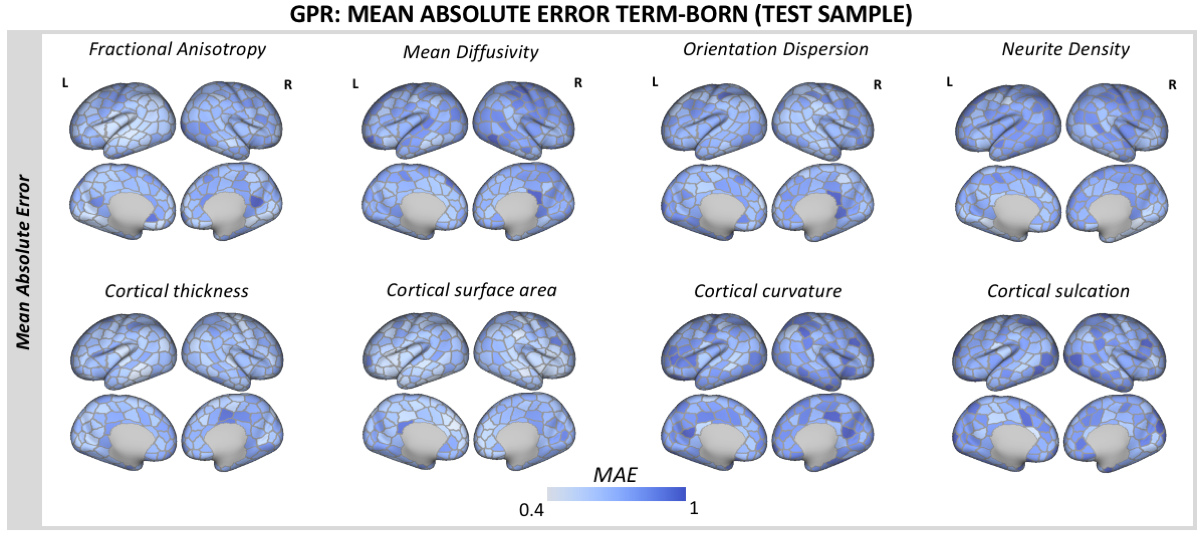

***Supplementary Figure 7.*** Mean absolute error surface maps (in units of standard deviation) derived from the GPR for the held-out term-born sample.

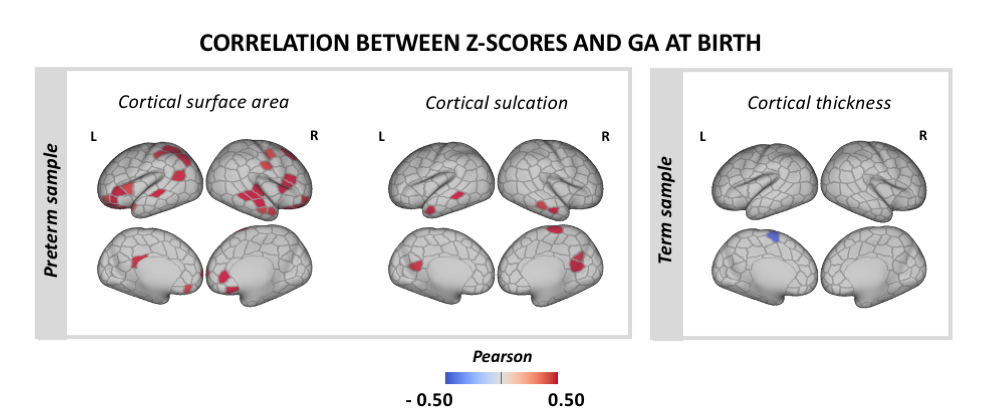

***Supplementary Figure 8.*** Association between extreme deviations from normative cortical development and GA at birth. Pearson’s correlation coefficients are shown only for parcels indicating significant correlation at p_mcfwe_< 0.05 in the preterm and term samples separately.

**Supplementary Tables:**

***Supplementary Table 1*.** Stages of quality control and exclusion criteria applied in sample size selection.

|  | **Term** | **Preterm** |
| --- | --- | --- |
| **Initial sample** | **400** | **109** |
| sMRI T_2_-weighted QC | -5 | -2 |
| dMRI QC score > 3.5 | -23 | -6 |
| dMRI cropped cortex | -76 | -10 |
| dMRI to sMRI misalignment | -2 | -5 |
| Surface QC | -18 | -10 |
| Incidental findings (clinical sign.) | -17 | * |
| **Final Sample** | **259** | **76*** |

*One preterm infant had a gross clinical abnormality and was excluded from the group-wise analyses. This infant’s data were kept in the individual GPR analyses.
